# Using 3D Invasion properties of RCC Cell Lines *In Vitro* to predict their Metastatic Potential *In Vivo*

**DOI:** 10.1101/2025.04.07.647527

**Authors:** B. Cesana, L. Nemoz-Billet, V. Azemard, C. Pillet, E. Bigot, N. Chaumontel, J.-L. Descotes, N. Osmani, J. G. Goetz, C. Cochet, O. Filhol

**Affiliations:** University Grenoble Alpes, Inserm, CEA, IRIG-Biosanté, UMR 1292, F-38000 Grenoble, France; Tumor Biomechanics Lab INSERM UMR_S1109 Strasbourg France, Université de Strasbourg France, Fédération de Médecine Translationnelle de Strasbourg (FMTS), Équipe Labellisée Ligue Contre le Cancer Strasbourg France; Centre hospitalier universitaire Grenoble Alpes, CS 10217, 38043 Grenoble cedex 9 France

**Keywords:** renal cell carcinoma, 3D models, patient-derived tumoroids, invasion assays, zebrafish embryos

## Abstract

Renal cell carcinoma (RCC) exhibits significant heterogeneity, making it challenging to predict tumor aggressiveness and therapeutic response. To improve prognostic accuracy and develop tailored treatment strategies, it is crucial to mimic both cancer cells and their microenvironment *in vitro*. Using a combination of *in vitro* and *in vivo* models, we investigated the invasive properties of three RCC cell lines—RCC10, RCC7 and 786-O— that displayed distinct signaling profiles, combining EMT characteristics and upregulation of key metastatic markers. Our findings revealed that RCC7 and 786-O exhibited greater metastatic potential than RCC10, as demonstrated by increased extravasation in zebrafish embryos and higher lung metastases in the chorioallantoic membrane (CAM) and mice models. Comparative pathway analysis indicated that RCC7 displays partial epithelial-mesenchymal transition (pEMT) characteristics and upregulates key metastatic markers. Furthermore, our 3D spheroid invasion model as well as our patient-derived RCC tumoroid system predicted accurately their metastatic behavior, closely mirroring their aggressiveness *in vivo*. Thus, these 3D models might be predictive of tumor outcome, underscoring their utility as reliable predictive tools for RCC progression and therapeutic response.

**Novelty and Impact:** Non-uniform distribution of genetic and phenotypic subpopulations within RCC tumors causes many tumors of similar histological grade to have vastly different metastatic potential. We show that 3D spheroids and RCC patient-derived tumoroid models more accurately reflect *in vivo* invasive behavior than traditional 2D assays, providing powerful predictive tools for RCC aggressiveness and metastatic disease. These findings have significant implications for precision oncology, enabling better preclinical evaluation of the metastatic risk to the patient.

## I. Introduction

Kidney cancer, particularly Renal Cell Carcinoma (RCC), accounts for approximately 2.4 % of global adult cancer cases, with clear cell RCC (ccRCC) being the most common subtype (70–80 % of cases) ^1^. Early-stage diagnosis has improved the five-year survival rate to 80 %, but 30% of RCC patients present metastatic RCC (mRCC) at diagnosis ^2^, reducing the five-year survival rate to under 10 % ^3^. Metastasis is a multi-step process in which cancer cells spread from a primary tumor to distant organs, leading to the formation of secondary tumors. This cascade involves several key steps: invasion of the nearby stroma, intravasation into blood or lymphatic vessels; vascular dissemination of circulating tumor cells (CTCs), extravasation into new tissues, and finally, colonization and metastatic growth ^4^. mRCC typically spread to the lungs (70 %), bones (32 %), lymph nodes (45 %), liver (18 %), and adrenal glands (10 %) ^5^, with 10 % of patients relapsing after surgery ^6^. Despite advances in targeted therapies and immunotherapy, mRCC prognosis remains poor, with a median overall survival of 10-15 months. The study of cancer aggressiveness relies on animal models to understand the mechanisms driving initiation, progression, and metastasis, with different *in vivo* models essential for analyzing each specific step. The chick chorioallantoic membrane (CAM) model provides a valuable platform that enables analysis of local invasion, intravasation, and metastasis in particular organs ^7^. The zebrafish model offers real-time imaging to study vascular dissemination and extravasation in an intact circulatory system ^8^. Finally, the mouse model remains the gold standard for studying metastatic colonization and metastatic growth ^9^. These models, and in particular PDX models (patient-derived xenografts) ^10^, have been pivotal in discovering and validating new cancer therapies in preclinical stages. However, the use of animals raises ethical concerns, pushing researchers to reduce their use. New approaches are emerging, such as organoids, which replicate human tissues and provide more relevant data without relying on animals ^11^. To analyze how tumor cells invade and interact within a more physiologically relevant context, the 3D cell cultures are embedded in hydrogels or other matrix systems that simulate the extracellular environment ^12^.

In our previous work, we developed a scaffold-free tumoroid model of ccRCC that preserves tumor self-organization, resident cell types, and gene expression patterns, closely mirroring patient tumors ^13^. However, it is crucial to confirm that their behavior in a 3D environment, such as an invasion-capable hydrogel, truly reflects the intrinsic aggressiveness of the original tumors. Although it is well-established that different RCC cell lines exhibit different mutational profiles, the relationship between their *in vivo* outcomes and *in vitro* behavior remains insufficiently characterized. In this work, we first evaluated the invasive potential of RCC cells *in vitro* using our 3D spheroid model and then compared their aggressive behavior using mouse xenografts, zebrafish embryos and CAM assays. We found that the aggressiveness evaluated by this 3D model correlates with the behavior in various *in vivo* models. Thus, this 3D model might be predictive of tumor outcome, thereby providing a functional system to screen for RCC targeted therapies.

## Material and methods

### Cell culture

786-O and ACHN cell lines (ATCC CRL-1932 and CRL1611) were maintained in RPMI-1640 medium, completed with 10% fetal bovine serum (FBS), 100 U/mL penicillin, and 100 µg/mL streptomycin. RCC10 cell line, provided by Gilles Pagès and Sandy Giuliano (University of Nice Sophia Antipolis), and RCC7 cell line, provided by Joel Le Maoult (St-Louis hospital/CEA Paris-Saclay) were cultured in complete DMEM (Gibco). RPTEC cell line was obtain from Evercyte and grown in ProXup media (Evercyte).

### Patients and Clinical Samples

Renal carcinoma samples were collected from patients who provided informed consent, with all procedures conducted in accordance with approval from the local ethics committee (Patient Protection Committee No. 2017-A0070251). The patients were enrolled as part of the Comborein clinical trial (NCT03572438). Fresh renal tumor tissues were obtained during partial or total nephrectomy surgeries performed for cancer treatment at the Urology Department of the University Hospital Center of Grenoble Alpes (CHUGA).

### Preparation of cell extracts and immunoblotting

RCC cell extracts were prepared as described in Supplementary Material and methods. Equal amounts of lysates (20 μg) were loaded onto 4-12 % gradient gels (Invitrogen) and summited to electrophoresis (NuPAGE buffer, 150 V, 1 h). Gels were transferred to PDVF membranes (100 V, 1 h) blocked (1 h, room temperature) with saturation buffer (5% BSA in TBST) and incubated with primary antibodies diluted in saturation buffer overnight. Immunoblotting was performed as described in supplementary Material and Method

### 2D cell assays

The proliferation, migration and invasion properties of 786-O, RCC10, and RCC7 cells were assessed as described in supplementary Material and Methods.

### Spheroid culture

RCC cells were counted and coated with magnetic nanoparticles (NanoShuttle, Greiner) at 1 µL per 40,000 cells through three centrifugation/resuspension cycles (1000 rpm - 5 min). Cells were then seeded (3000 cells/well) in a 96-well U-bottom plate pre-coated with 20 mg/mL poly (2-hydroxyethyl methacrylate) (Sigma-Aldrich). The plate was incubated at 37°C, 5% CO_2_ for 3 days on a Spheroid-Drive support (Greiner) to form mature spheroids ^13^.

### Tumoroid culture

Generation of tumoroids was performed as previously described ^13^. Briefly, fresh patient tumor samples were minced, enzymatically digested, and mechanically processed with the GentleMACS Dissociator. Cells were coated with magnetic NanoShuttle (Greiner, 1 µL/40,000 cells) before being seeded (30,000 cells/well) in flat-bottom, low-attachment 96-well plates (Greiner). Human tumoroids were cultured in TumorMACS® medium (Miltenyi Biotec) supplemented with 10% FBS (GIBCO) and 100 U/mL penicillin-streptomycin. Plates were placed on the spheroid-drive support and incubated for 21 days at 37°C, 5% CO^2^.

### 3D cell invasion

Mature spheroids/tumoroids were embedded in a Collagen I Rat (3mg/ml) + Fibronectin (10 µg/ml) hydrogel in low-attachment 96-well plate. After polymerization, DMEM-F12 was added and cultures were maintained for 7 days. For endpoint invasion analysis, spheroids/tumoroids were stained (Hoechst 33342, Live/Dead viability kit Invitrogen) and imaged (AxioObserver Z1). ImageJ was used to measure invasion distance, area, and cell count. For time-lapse analysis, spheroids/tumoroids were monitored in CELLCYTE X™ equipped with the “3D Spheroids” module (4X, hourly), with invasion speed and cell tracking analyzed via ImageJ. Immunofluorescent tumoroid labeling was carried out as described in supplementary Material and Methods.

### Animal studies

All animal studies were approved by the institutional guidelines and those formulated by the European Community (EU Directive 2010/63/EU) for the Use of Experimental Animals.

### Mice

786-O, RCC10 and RCC7 adherent cells were transduced with Lenti-II-CMV-Luc-IRES-GFP virus (ABM, Germany) at a multiplicity of infection of 1–5 with 8 μg/mL polybrene (Sigma). Cells (1 × 10^6^) were injected into the retro-orbital sinus of athymic nude mice (6 weeks-old female BALB/c Nude, CByJ.Cg-Foxn1nu 522 /J; Charles River). Tumor growth was followed by intraperitoneal injection of 150 mg/kg D-luciferin Potassium salt using IVIS Imager (Perkin Elmer). Once tumors reached a critical size, mice were injected with luciferin before being euthanized. Lungs were inflated with luciferin via tracheal injection. Primary metastatic organs (kidneys, lungs, liver, femur) were dissected, imaged with an IVIS Imager, and fixed in 4% paraformaldehyde (2 h). (Mice facility A1-0741, #9436-2017032916298306).

### Zebrafish embryos

Zebrafish experiments were performed as previously described ^14^. Briefly, Tg(fli:gfp) zebrafish (Danio rerio) embryos were housed at 28°C in a 0.3X Danieau solution. At 48 h post-fertilization (hpf), embryos were immobilized in a low-melting point 0.8 % agarose pad containing 650 mM tricaine to allow stable injection conditions. Using a Nanoject II microinjector (Drummond) and custom-microforged glass capillaries (internal diameter 25–30 µm), 13.8 nL of RCC cell suspension (100 × 10^6^ cells/mL) was introduced into the duct of Cuvier of each embryo under a M205 FA stereomicroscope (Leica). The 786-O, RCC10 and RCC7 cells used in this study had been previously transduced with an H2B-mCherry lentivirus (Plasmid #20972), resulting in red fluorescent labeling of their nuclei. Images of the caudal plexus were acquired at 3, 24 and 48 h post-injection (hpi) for each embryo with an inverted Olympus IX83 with a CSU-W1 spinning disk head (Yokogawa), an ORCA Fusion Digital CMOS camera (Hamamatsu), and an UPL SAPO 30x 1.05 NA silicone immersion objective. All zebrafish procedures were performed in accordance with French and European Union animal welfare guidelines and supervised by local ethics committee (Zebrafish facility A6748233; APAFIS 2018092515234191).

### Chicken embryos

Fertilized White Leghorn eggs were sourced from Couvoir Hubert, France. On embryonic development day 9 (EDD9), a 50 µL volume of cell suspension containing 3 × 10^6^ cells (786-O, RCC10, or RCC7) was then grafted onto the CAM. Tumor growth and embryo viability was monitored regularly. On EDD18, tumors were excised, washed with PBS and weighed. Lungs and lower CAM were collected, snap frozen and stored at −80°C. In compliance with French regulations, ethical approval was not necessary for experiments involving oviparous embryos (Decree No. 2013-118, February 1, 2013; Article R-214-88).

### Immunohistochemistry

Immunochemistry was performed on a sample of formalin-fixed paraffin-embedded tissue sections. Briefly, formalin-fixed paraffin-embedded 4 μm thick sections were first deparaffinized in xylene and rehydrated in alcohol. Slices were counterstained for 20 s with pure Harris Hematoxylin (Sigma) followed by washing using water and stain for 3 min using Eosin (Sigma). Slices were then dehydrated in ethanol 100 % for 2 × 5 min and with Xylene for 2 × 2 min, mounted using Merkoglass (Sigma) and visualized under microscope Zeiss™ AxioVision Rel. Software 4.X.

### qPCR

Total nucleic acid was extracted using the NucleoSpin 96 Tissue kit (Macherey–Nagel). Quantitative PCR (qPCR) was performed on human-specific Alu sequences to assess the relative quantity of human DNA in the lower CAM and the lungs of chicken embryos. qPCR was performed using SsoAdvanced Universal Probes Supermix (Bio-Rad) in a CFX96 Real-Time System (Bio-Rad) following the manufacturer’s protocol. We used Alu sequences primers (Custom PrimePCR Assay, Bio-Rad) based on the protocol described by Funakoshi et al ^15^, summarized in Figure S1C. The probe was labeled with the fluorescent dye 6-carboxy-fluorescein (6-FAM). qPCR was conducted under the following conditions: an initial cycle at 95°C for 3 minutes, followed by 45 cycles at 95°C for 15 seconds, 56°C for 30 seconds. Data analysis was conducted utilizing the CFX Manager Software V3.1 (Bio-rad).

### Statistical analysis

GraphPad Prism 10.2.1 was used for the generation of graphs and statistical analysis. Significance was determined using a one-way ANOVA with Turkey’s multiple comparisons test.

## II. Results

### Comparative Analysis of RCC Cell Lines using 2D cell culture Methods

In this study, we selected three different renal cancer cell lines: RCC10, 786-O and RCC7, each derived from patients with RCC (as indicated in Table 1, (a)^16^, (b)^17^, (c)^18^). These cell lines exhibit distinct mutational profiles, particularly regarding the VHL status, which regulates HIF-α degradation and drives tumorigenesis when mutated ^19^. In order to functionally investigate the pre-existing heterogeneity of these RCC cell lines, we compared their proliferation, migration and invasion properties using conventional 2D techniques. Since cell proliferation is critical to support both primary tumor and metastasis growth, we quantified the proliferation rate of each cell line over 60 h. Tracking of individual cells and measurement of cell proliferation rate in % over time for each cell line using the CELLCYTE X™ software gave the following average rate of proliferation: RCC7 (1.494 %/h), 786-O (1.313 %/h) and RCC10 (1.256 %/h) (Figure S1 A-B).

**Table 1:** Known characteristics of three human ccRCC cell lines used in this study. U.S. means unspecified.

| Cell line | Age | Sex | Origin | Tumor Type | Size in suspension | Doubling Time | VHL mutation | Other mutations | Ref |
| --- | --- | --- | --- | --- | --- | --- | --- | --- | --- |
| **RCC10** | U.S. | U.S. | U.S. | ccRCC | 15 µm | 29h | VHL - | U.S. | a |
| **786-O** | 58 | Male | Caucasian | ccRCC | 15 µm | 24h | VHL - | TP53, PTEN, TERT | b |
| **RCC7** | U.S. | U.S. | U.S. | ccRCC | 15 µm | 24h | VHL + | U.S. | c |

In contrast, the migration rate of each cell line determined in wound-healing assays (Figure S1 C-D) gave the following average velocity of wound closure: RCC7 (0.669 %/h), RCC10 (1.221 %/h) and 786-O (1.334 %/h) (Figure S1 E). During tumor expansion, cancer cells must invade the extracellular matrix and surrounding tissues, a process that relies on the degradation and proteolysis of ECM components ^20^. The invasiveness of each cell line evaluated with a Matrigel-coated Boyden chamber assay showed that RCC10 cells exhibited a significantly lower invasive potential compared to the other two RCC cell lines (Figure S1 F-G).

To investigate the molecular mechanisms underlying these differences in invasiveness, migration and proliferation, we analyzed some key signaling pathways in RCC10, 786-O, and RCC7, alongside ACHN ^21^ and normal renal epithelial cells (RPTEC ^22^) using western blotting (Figure S2A). Consistent with their VHL status (Table 1), RPTEC, ACHN, and RCC7 cells did not express the hypoxia-inducible factor HIF-2α, whereas RCC10 and 786-O cells did. Analysis of Epithelial-to-Mesenchymal Transition (EMT) markers, revealed reduced E-cadherin and elevated N-cadherin and vimentin expression in all RCC cells compared to RPTEC, a switch representing a hallmark of EMT ^23^. In accordance, transcription factors such as Snail1 and ZEB2, activating EMT-promoting signaling pathways, were also overexpressed in all RCC cells. However, RCC7 cells displayed lower EMT markers expression reminiscent of a partial EMT-like phenotype (p-EMT), a state associated with increased plasticity and invasive potential, enabling cancer cells to retain some epithelial characteristics while acquiring mesenchymal traits ^24, 25^.

β1 integrin analysis revealed a single band in RPTEC and ACHN cells, while RCC cells displayed shifted migrating bands, suggesting altered glycosylation. Given that our monoclonal antibody targets the NH2-terminal ectodomain, these variations likely reflect N-glycosylation modifications, which have been linked to β1 integrin activation and metastatic progression ^26^.

Additionally, matrix metalloproteinase 2 (MMP2), a critical enzyme involved in ECM degradation, was significantly upregulated in RCC7 cells. Interestingly, it has been reported that MMP2 is involved in multiple steps of the metastatic cascade playing an important role in EMT and cell dissemination ^27^. The adhesion properties of RCC cells are also influenced by the chemokine receptor CXCR4 ^28^, which was highly expressed in RCC10 and 786-O cells but nearly undetectable in RCC7 cells. This observation aligns with reports that metastatic cancer cells tend to exhibit lower CXCR4 expression compared to primary tumors ^29^. Moreover, RCC7 cells exhibited lower levels of OCT4, a stemness marker associated with dedifferentiation and RCC tumor initiation ^30^, suggesting that they are in a more differentiated state compared to the other RCC cells. Programmed death-ligand 1 (PD-L1), an immunosuppressive protein that inactivates T cells by binding to the inhibitory receptor programmed death-1 (PD-1), was strongly expressed in RCC7 cells as compared to the other RCC cells. In tumors, this immune checkpoint system is exploited to evade immune surveillance, a process known as immune evasion. Interestingly, PD-L1 expression is now established as a strong predictor of unfavorable prognosis in RCC ^31^.

Regarding changes in phosphorylation in signaling pathways (Figure S2B), AKT phosphorylation was upregulated in RCC10 and 786-O but impaired in RCC7, consistent with studies linking AKT inhibition to increased invasiveness ^32^. Paxillin1 phosphorylation, crucial for EMT and migration, was reduced in RCC10 and RCC7, mirroring findings from zebrafish models of single-cell migration ^33^. Overall, RCC7 cells exhibit a distinct landscape of signaling pathways, combining p-EMT characteristics with altered phosphorylation patterns, which may contribute to their exacerbated invasive behavior.

Given the limitations of 2D cell cultures in accurately evaluating cell aggressiveness, we hypothesized that our 3D spheroid model ^13^ could generate phenotypically diverse spheroids based on their distinct signaling features.

### Comparative analysis of RCC cell invasion in a 3D environment

To study cell invasion in a 3D context, spheroids made from RCC cells (786-O, RCC7 and RCC10) were embedded in an hydrogel, mimicking the mechanical properties of tumor tissue ^34^. We found that the 4.864 ± 0.013 kPa hydrogel’s stiffness measured via texturometry, aligns closely with both the human kidney cortex and medulla (4–7 kPa) and tumor-adjacent tissues (2–6 kPa) ^35^, supporting its relevance in the study of a renal cancer model. Over the course of 7 days, RCC10 spheroids remained compact within a main cell-packed body, whereas in 786-O and RCC7 spheroids, cancer cells escaped, invaded, and survived within the surrounding 3D matrix (Figure 1A). RCC7 cells were found to be the most invasive, showing a significant higher maximum invasion length (Figure 1B), invasion area (Figure 1C), and number of invasive cells (Figure 1D), while 786-O cells displayed significant but lower invasiveness and RCC10 very low invasive properties. We recorded invasive cells over a 7-day period (Supplementary Video 1-3) and calculated the average invasion speed for each cell line: RCC7 (10.55 µm/h), 786-O (5.73 µm/h) and RCC10 (0.81 µm/h) (Figure 1E, 1F). Tracking the trajectory of single invading cells revealed distinct migratory behaviors (Figure 1G). RCC10 cells exhibited a directed, forward-biased migration pattern, moving mostly in one direction with occasional zigzagging. In contrast, RCC7 and 786-O cells showed more erratic, non-linear trajectories, with frequent directional changes, pauses, and reversals, suggesting a more exploratory motility pattern.

**Fig 1:**
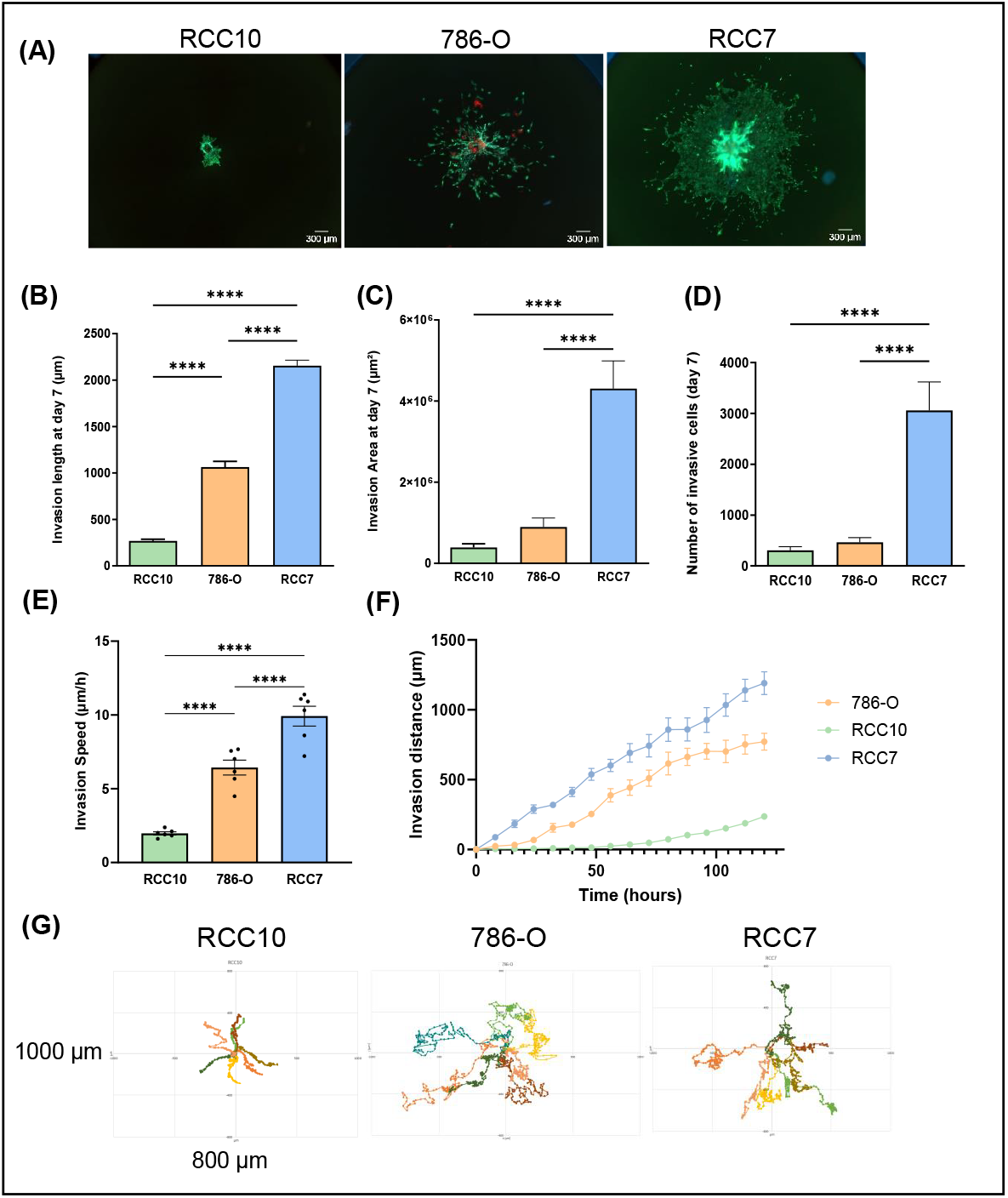
3D spheroid invasion assay. **(A)** Representative images of RCC7, 786-O and RCC10 spheroids cultured in Collagen I and Fibronectin hydrogel visualized after 7 days with Live and Dead staining (Calcein staining living cells in green, Hoechst 33342 staining nuclei in blue and Ethidium Homodimer I staining dead cells in red). Scale bar = 300 µm. **(B)** Maximum invasion length of RCC cell lines invading through hydrogel after 7 days of culture. **(C)** Average invasion area of ccRCC cell lines invading through hydrogel after 7 days of culture. **(D)** Number of cells invading the hydrogel after 7 days of culture. **(E)** Invasion speed of each cell line. **(F)** Invasion speed of RCC7, 786-O and RCC10 spheroids embedded in hydrogel measured during 5 days. **(G)** Representative rose plot of the invasion of cells through hydrogel. Data show mean ± SEM, RCC7 n = 10, 786-O n = 16, RCC10 n = 15, ACHN n = 12. Significance was assessed using a one-way ANOVA with Tukey’s multiple comparison test comparing each cell line to every other cell line. (\**p*<0.05;**** *p*<0.0001)

Although *in vitro* analysis of RCC cell lines in this 3D invasion assay highlighted their sharp differences in invasiveness, it was essential to determine how these observations translate to *in vivo* tumor aggressiveness and metastatic potential.

### In vivo analysis of RCC cells in the chicken egg chorioallantoic membrane (CAM) model

To study the capacities of RCC cells to form a primary tumor and intravasate, we employed the CAM model. This model is particularly valuable in cancer research as it provides a highly vascularized, immunodeficient environment that supports rapid tumor growth, angiogenesis, and spontaneous metastasis, making it an excellent system for studying tumor progression *in vivo*. It also allows for the assessment of tumor cell graft toxicity by monitoring embryo survival rates, providing insights into the potential impact of tumor burden on host viability. RCC cells were grafted in the upper CAM on embryonic development day 9 (EDD9). On EDD18, samples from each specimen were collected and analyzed (Figure 2A). All three cell lines successfully formed tumors *in ovo*. Morphologically, RCC7 tumors were strongly hemorrhagic, RCC10 tumors displayed a glandular structure, and 786-O tumors were largely avascular (Figure 2B). Tumors from RCC10 and 786-O were uniform in size, while RCC7 tumors were more heterogeneous and the largest, averaging 23.8 mg, compared to 8.1 mg for 786-O and 5.8 mg for RCC10 (Figures 2C and 2D). Metastasis was quantified using sensitive qPCR for human Alu sequences in genomic DNA from the lungs or lower CAM (Figure S3A-C). All three cell lines formed metastatic foci in the lower CAM, though variability was observed, showing no metastases with one egg each in the 786-O and RCC7 groups and five in the RCC10 group. Similarly, metastatic foci in the lungs were detected in 2/8 eggs for 786-O and RCC10, and 3/8 for RCC7 (Figure 2E). Notably, no significant signs of embryo toxicity were observed following grafting with any of the cell lines, although a slightly lower survival rate was noted in the RCC7-grafted group (Figure S3D). This model can effectively highlight distinct metastatic capacities between the RCC cell lines.

**Fig 2:**
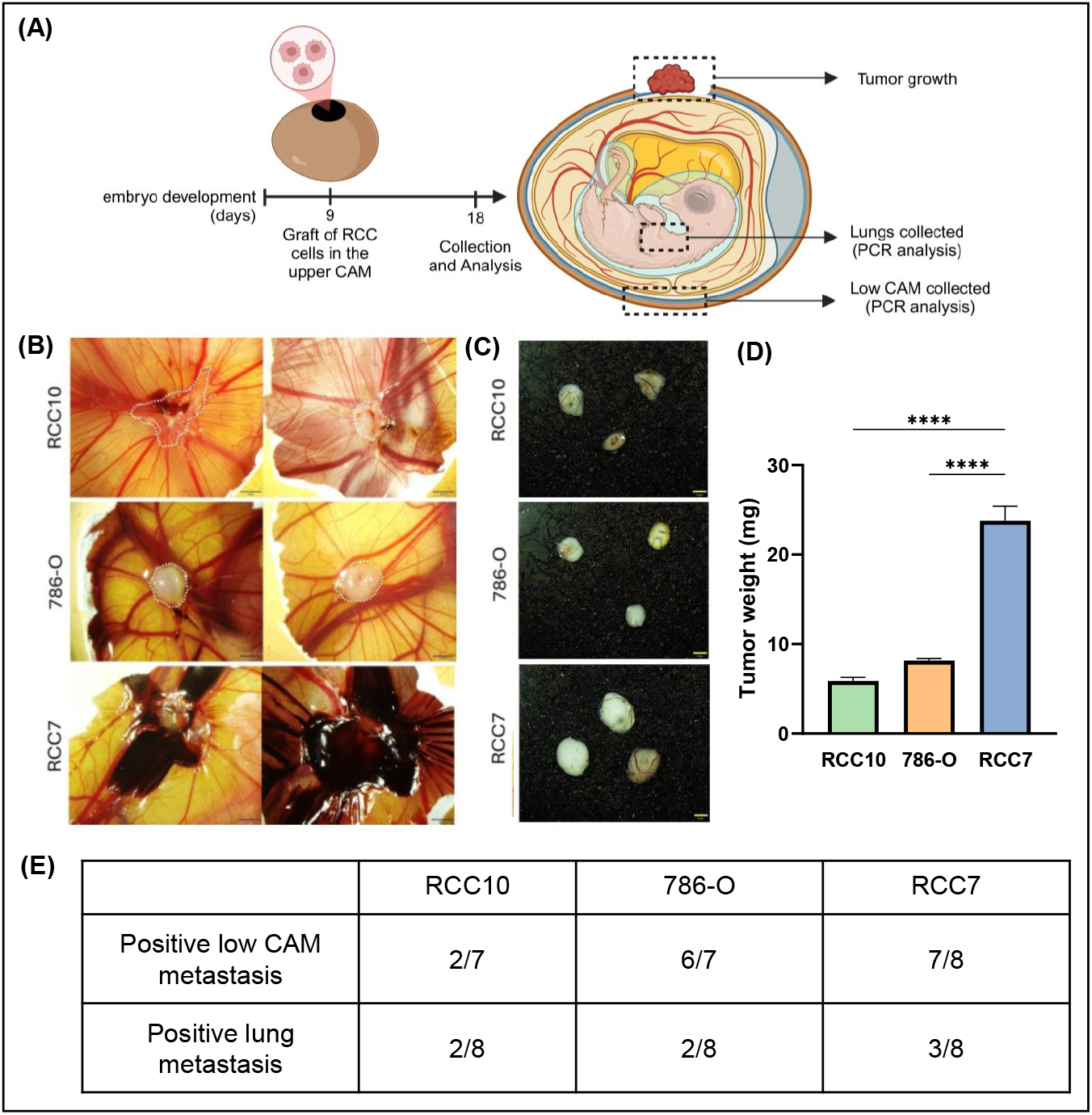
RCC cells have different tumor growth and metastatic potential in the Chick embryo chorio-allantoic membrane (CAM) model. **(A)** Scheme of the experimental approach of RCC cells injection into the Chorio-Allantoic Membrane of chick embryos. **(B)** Representative images of tumor derived from RCC10, 786-O and RCC7 cells in ovo at E16. For RCC7 tumors, the important accumulation of blood prevents to distinctly locate the tumor. **(C)** Illustrative pictures of RCC10, 786-O and RCC7-induced tumors (post-collection). **(D)** Mean tumor weight (mg) measured in the different experimental groups at the end of the study. **(E)** Number of embryos that showed a positive signal in the analysis of the metastatic invasion, measured by qPCR for human specific Alu sequences in the lower CAM and in the lungs. Data are shown as mean with SEM with n = 8 animals/group and n = 3 for the control. Significance was assessed using a one-way ANOVA with Tukey’s multiple comparison test comparing each cell line to every other cell line. (**** *p*<0.001).

### In vivo extravasation of RCC cells in zebrafish embryos

The zebrafish model has emerged as a powerful system for studying the dynamic interactions between cancer cells and the vasculature, providing high-resolution, real-time imaging of circulating tumor cells within an intact circulatory system ^36^. Unlike the CAM model, which allows for the assessment of tumor growth and metastasis, the zebrafish model enables direct visualization of key metastatic events, including intravascular arrest, adhesion, and extravasation, over a short timescale. To investigate RCC cell line behavior, we injected mCherry-labeled RCC7, 786-O, and RCC10 cells, into the blood circulation of zebrafish embryos, monitoring their adhesion and extravasation in the caudal plexus at 3-, 24-, and 48-hours post-injection (hpi). The caudal plexus, a well-characterized vascular network in the posterior tail region, serves as an accessible and physiologically relevant model for studying blood flow dynamics, tumor cell adhesion, and extravasation, making it highly valuable for cancer and vascular research. This network comprises multiple vessels, including dorsal aorta (DA), arterio-venous junction (AVJ), caudal vein (CV), providing distinct routes for circulating cells (Figure 3A). At 3 hpi, no significant differences in intravascular arrest were observed, confirming that an equivalent number of cells were initially injected (Figure 3B). By 24 hpi, RCC7 and 786-O cells exhibited greater adhesion than RCC10 (Figure 3C). Representative images illustrate cells arrested at this time point (Figure 3D), while heatmaps quantify the number and spatial distribution of stably arrested CTCs in the caudal plexus (Figure 3E) showing a preference for the CV especially for RCC7 and 786-O (Figure 3F). At 48 hpi, RCC7 and 786-O remained more abundant in the caudal plexus, suggesting enhanced survival or adhesion compared to RCC10 (Figure 3G). The zebrafish model uniquely enables the quantitative assessment of extravasation at the single-cell level, providing high-resolution insights that surpass those obtainable with bioluminescence imaging in mouse or CAM models. To ensure accurate identification of extravasated cells, we employed orthogonal views to confirm their localization outside the vessel boundaries (Figure 3H). Interestingly, 786-O and RCC7 cells exhibited higher extravasation rates than RCC10 (Figure 3I-J), highlighting potential differences in metastatic potential among these RCC cell lines.

**Fig 3:**
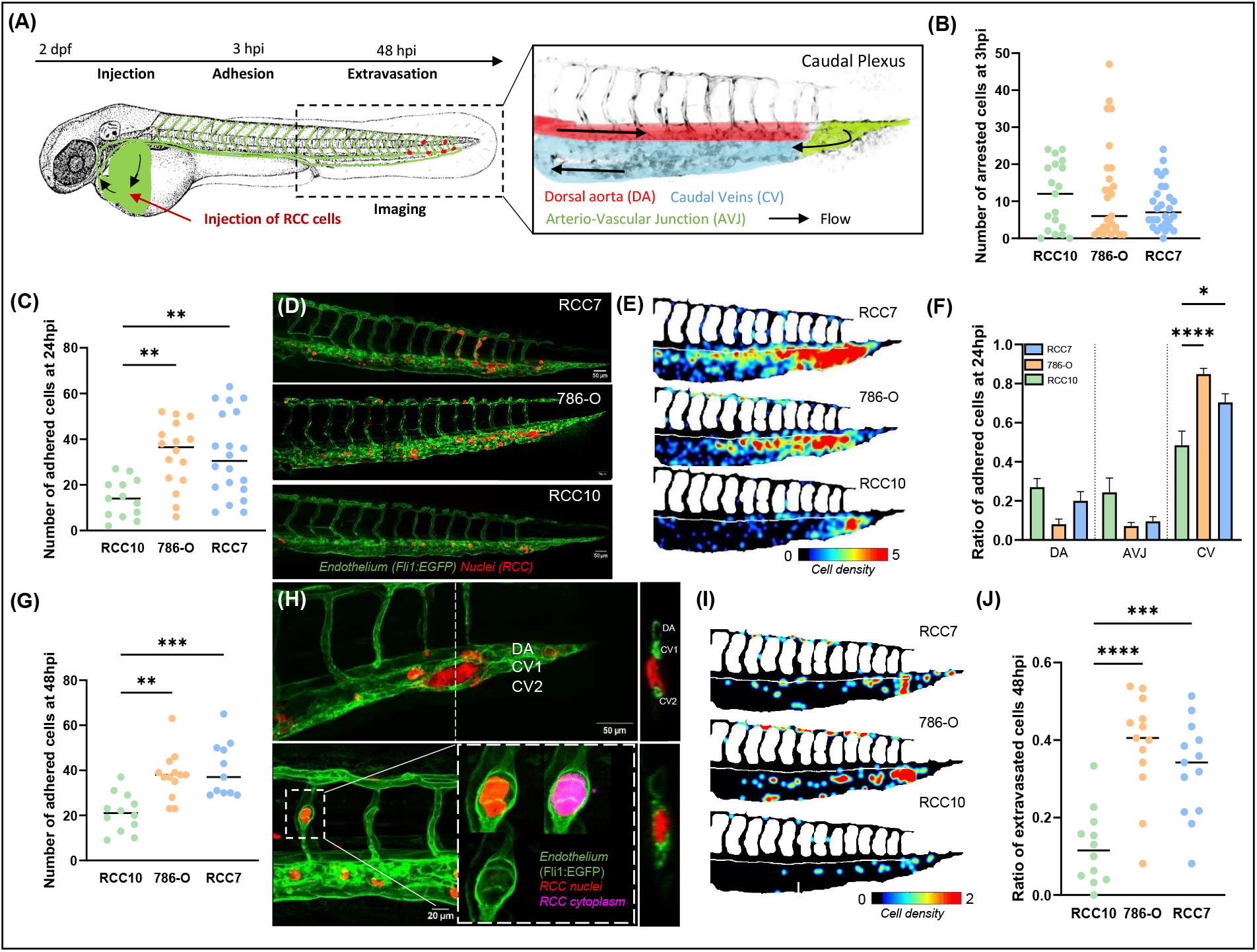
Adhesion and extravasation capacities of RCC cells in zebrafish in vivo model. **(A)** Scheme of the experimental approach using 2 dpf Tg(Fli1:EGFP) zebrafish embryos as a metastatic cascade model. Right panel: blood flow in the caudal plexus regions: dorsal aorta (DA), arterio-venous junction (AVJ), caudal veins (CVs). The adhesion pattern of RCC7, 786-O or RCC10 cells was imaged 3, 24 and 48 hours post-injection (hpi). **(B)** Total number of arrested cells at 3 hpi. The graph shows the mean ± SEM of 2 independent experiments (RCC10 n = 19; 786-O = 27; RCC7 n = 28). **(C)** Total number of arrested cells at 24 hpi **(D)** Representative images of cells arrested at 24 hpi. **(E)** The heatmaps show quantification of the number and location of arrested RCC cells at 24 hpi in the caudal plexus. **(F)** Ratio of cells adhered at 24 hpi over the total number of cells and their localization (see A) of cells adhered at 24 hpi were measured. The graphs shows the mean ± SEM of 2 independent experiments (RCC10 n = 13; 786-O = 16; RCC7 n = 20). **(G)** Total number of arrested cells at 48 hpi. The graph shows the mean ± SEM (RCC10 n = 12; 786-O n = 13; RCC7 n = 11). **(H)** Representative images (left) and its orthoslice (right) of extravascular cells at 48 hpi. **(I)** The heatmaps show quantification of the number and location of extravascular RCC cells at 48 hpi in the caudal plexus. **(J)** Ratio of cells extravasated over the total number of cells was measured at 48 hpi Significance was assessed using a one-way ANOVA with Tukey’s multiple comparison test comparing each cell line to every other cell line. (\**p*<0.05; ** *p*<0.01; *** *p*<0.001; **** *p*<0.0001).

### In Vivo Characterization of Metastasis in RCC cell-derived Xenografts

While the zebrafish and CAM models provide valuable insights into the early stages of metastasis, including tumor growth, intravascular arrest, adhesion, and extravasation, the mouse model remains indispensable for studying metastatic colonization and growth in a fully developed mammalian microenvironment. This model enables the evaluation of late-stage metastasis and organ-specific tumor progression, offering crucial insights that cannot be captured in avian or fish models. To compare the metastasizing capacities and the aggressiveness of the three cell lines, we injected cultured cells retro-orbitally into immunocompromised mice, enabling systemic dissemination and facilitating the evaluation of organ-specific colonization and tumor spread. To monitor tumor growth, cells were previously transfected with a luciferase plasmid, allowing for real-time tracking of disease progression *in vivo* using bioluminescence imaging (Figure 4A). At the end of the study, specific organs were analyzed, including the lungs, liver, femoral bone and kidneys, revealing metastatic dissemination exclusively in the lungs. All mice injected with RCC7 or 786-O cells developed lung metastases, while no metastasis was detected in the RCC10-injected group (Figure 4B-C). Interestingly, RCC7-injected mice developed metastases remarkably quickly, with detectable lung lesions emerging as early as 4 weeks post-injection, compared to 11 weeks for 786-O-injected mice. In contrast, mice injected with RCC10 cells showed no evidence of metastasis by the planned 12-week endpoint, underscoring significant differences in metastatic progression among the different cell lines (Figure 4D-E). Thus, these data revealed that these RCC cell lines, while having the same behavior in 2D cell culture, exhibit differential metastatic capacities.

**Fig 4:**
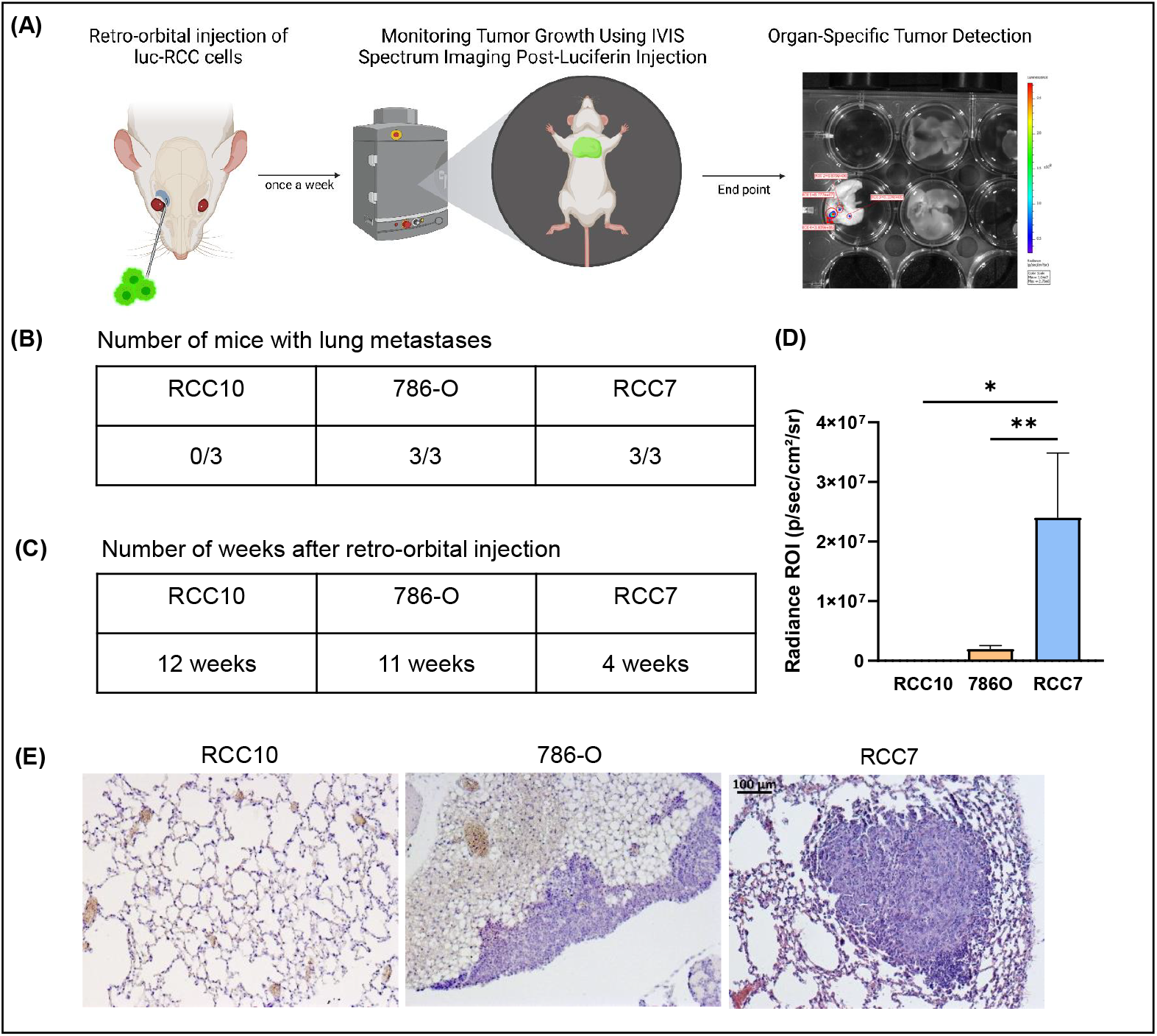
Metastatic Potential of RCC Cells in Murine Lungs. **(A)** Scheme of the experimental approach of retro-orbital injection of luciferase-transduced RCC cells into BALB/c Nude female mice. Lungs, liver, femoral bone and kidney were specifically screened for metastasis. **(B)** Number of mice that developed lung metastases for each cell lines. **(C)** Time of metastasis onset after retro-orbital injection. **(D)** Average radiance intensity measured after luciferin injection (200µl, 15 mg/ml) with IVIS® Spectrum. **(E)** Representative tissue sections stained with H&E, obtained from lungs of mice after retro-orbital injection of RCC cells. Scale bar = 100 µm. Data are shown as mean with SEM with n=3 animals/group. Significance was assessed using a one-way ANOVA with Tukey’s multiple comparison test comparing each cell line to every other cell line. (\**p*<0.05; ** *p*<0.01).

### Comparative analysis of fresh patient-derived tumoroid invasion in a 3D environment

To determine whether the invasive properties of renal cancers are the primary drivers of metastatic disease in human patients, we tested tumoroids derived from two RCC patients disclosing different ISUP grading ^37^, MC-156 (T3A, ISUP 3) and PD-157 (T2A, ISUP 2), using the same invasion model. Upon receipt, tissues were digested to a single-cell suspension, and processed to generate ccRCC tumoroids that provide a more clinically relevant representation of tumor heterogeneity, potentially capturing patient-specific invasive behaviors not observed in established cell lines. We observed significantly different tumoroid invasion behaviors (Figure 5A): MC-156 tumoroids exhibited high invasion with a maximum length and average speed of 5.02 µm/h, compared to 0.90 µm/h for PD-157 tumoroids (Figure 5B-D). MC-156 cells also displayed an exploratory motility pattern, with dynamic and extensive movement, while PD-157 cells were more restrained (Figure 5E). Immunostaining with the ccRCC marker CAIX ^38^ confirmed that most cells invading from MC-156 tumoroid were tumor-derived renal cells (Figure 5F). Although preliminary, these data suggest that this 3D model can be used to measure the invasive capacity of ccRCC tumor biopsies. Thus, combining histopathological assessment with 3D invasion traits may better correlate to clinical outcomes and serve as well as a valuable tool for screening RCC-targeted therapies.

**Fig 5:**
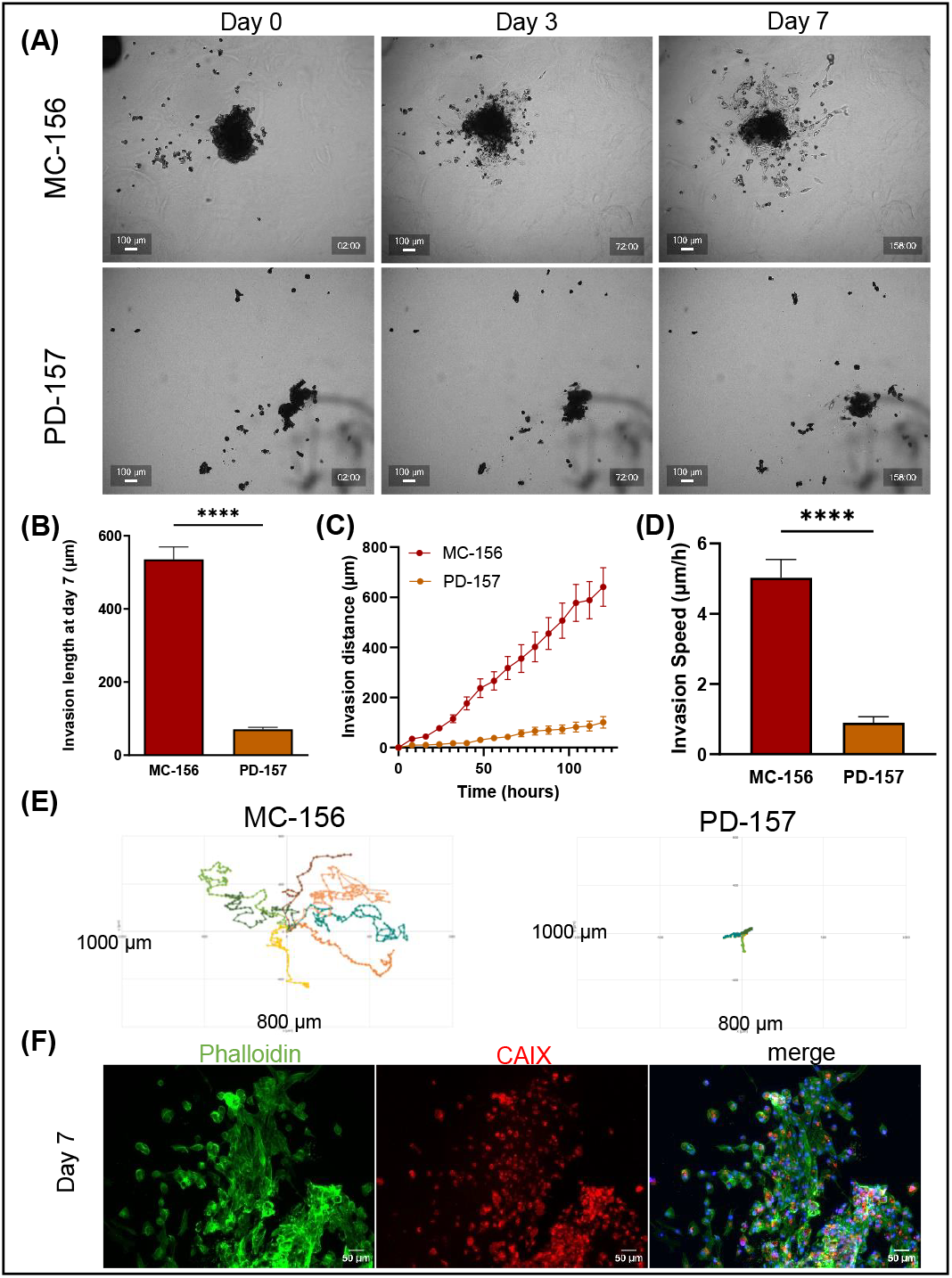
3D human renal tumoroid invasion assay. **(A)** Representative images of MC-156 and PD-157 tumoroids cultured in Collagen I and Fibronectin hydrogel, at 0, 3 and 7 days of culture. (Scale bar = 100 µm). **(B)** Maximum invasion length of RCC cells invading through hydrogel after 7 days of culture. **(C)** Average invasion area of RCC cells invading through hydrogel after 7 days of culture. **(D)** Invasion speed of RCC7, 786-O and RCC10 spheroids embedded in hydrogel measured during 5 days. **(E)** Representative rose plot of the invasion of tumoroid cells through hydrogel. Data show mean ± SEM, MC-156 n = 12, PD-157 n = 12. **(F)** Immunostaining of MC-156 invading cells (CAIX antibody staining renal cells in red, phalloidin staining F-actin in green and Hoechst staining nuclei in blue). Significance was assessed using a one-way ANOVA with Tukey’s multiple comparison test comparing each cell line to every other cell line. **** *p*<0.0001).

## III. Discussion

In this study, we aimed to characterize the metastatic potential of three distinct RCC cell lines—RCC10, RCC7 and 786-O—using complementary *in vitro* and *in vivo* models. Our findings revealed that distinct metastatic behaviors evaluated through an *in vitro* 3D model and *in vivo* studies may be associated with significant genetic and phenotypic heterogeneity among these RCC cell lines.

To evaluate tumor growth and metastatic progression of these RCC cell lines, we employed three animal models, each offering distinct advantages : 1) The CAM model allows investigating tumor growth, angiogenesis, local invasion and metastasis detection; 2) The zebrafish model provides visualization of CTC and extravasation within an intact circulatory system ^8^; 3) The mouse model enables the assessment of tumor aggressiveness with systemic dissemination and organ-specific metastasis.

The three RCC cell lines successfully engrafted in the CAM model. However, RCC7 tumors were larger, hemorrhagic and exhibited greater heterogeneity. Moreover, the three cell lines displayed uneven metastatic spread both in the lower CAM and in lungs. Nevertheless, this analysis alone did not allow to statistically evaluating their respective aggressiveness.

To further investigate metastatic events, particularly extravasation, we employed zebrafish embryos ^14^. This model allowed us to assess the adhesion and extravasation capacities of RCC7, 786-O, and RCC10 cells in a dynamic environment. All three cell lines displayed comparable initial vascular arrest, suggesting similar levels of circulation and entrapment. However, by 24 hpi, RCC7 and 786-O cells remained more abundant in the caudal plexus compared to RCC10, indicating enhanced survival or potential occlusion capabilities within the circulatory environment ^8^. By 48 hpi, RCC7 and 786-O cells exhibited significant higher extravasation rates compared to the RCC10, reinforcing their superior ability to traverse the endothelial barrier. Interestingly, these findings align with prior work by Osmani *et al*., who demonstrated that the highly metastatic D2A1 cell line extravasates more efficiently than its less aggressive counterpart (D20R) in zebrafish embryos ^39^. This further supports the notion that extravasation efficiency is closely linked to metastatic potential, underscoring the relevance of this model for studying tumor cell dissemination.

Finally, in the mouse model, RCC7 and 786-O cells also formed lung metastases, whereas RCC10 cells did not, confirming their different metastatic behavior in accordance with the other two *in vivo* models. RCC7 cells exhibited a faster metastatic onset, suggesting that these cells may possess higher metastatic aggressiveness. Altogether, the rapid metastasis onset of RCC7 cells and larger lesions in mice may be correlated with both their metastatic capacity in chicken embryos and their higher adhesive potential in zebrafish vessels. In contrast, the limited extravasation capacities of RCC10 cells in zebrafish maybe linked with their inability or lower capability to form metastases in mice and in CAM model respectively.

Despite their advantages, *in vivo* models present ethical, financial, and logistical challenges, making *in vitro* methods still essential for initial cancer research investigations.

Since we suspected that distinct metastatic behaviors among these RCC cell lines may be associated with significant genetic and phenotypic heterogeneity, we performed a comparative analysis of key signaling pathways. In this analysis, RCC7 cells exhibit lower expression of EMT markers (Snail1, N-cadherin, vimentin), suggesting partial EMT traits with some regained epithelial identity. It has been reported that tumor cells can reactivate plasticity programs to modulate differentiation processes, including EMT and mesenchymal-to-epithelial (MET) transitions, key mechanisms in carcinogenesis ^40^. Previous studies have identified a pre-existing EMT continuum in cancers, comprising epithelial, intermediate EMT, quasi-mesenchymal, and fully mesenchymal states contributing to tumor heterogeneity ^40^. The signaling pathway profile of RCC7 cells aligns with previous evidence that EMT plasticity contributes significantly to metastatic potential ^25^. Additionally, RCC7 displayed distinct CXCR4 and OCT4 patterns, indicating a more differentiated state than RCC10 and 786-O. Notably, RCC7 also overexpressed MMP2, an enzyme critical for metastatic progression ^41, 42^. Furthermore, the strong expression of PD-L1 observed in RCC7 cells may be associated with their aggressiveness, protecting them from cytotoxic T cells, thereby potentiating a high immune-suppressive effect ^31, 43^. Alternatively, this ligand was reported to have an intrinsic role in cancer cell migration and invasion independently of its binding to PD-1 receptors on T cells ^44^.

Common *in vitro* assays, such as proliferation, migration, and invasion assays, are widely used to characterize the metastatic potential of cancer cells ^45^. However, in our study, these 2D assays failed to confirm our *in vivo* experiments, underscoring the need for more physiologically relevant models. To address this, we employed a 3D spheroid invasion model, which better mimics *in vivo* tumor behaviors and reflects the intrinsic heterogeneity of metastatic capacities ^46^. We embedded RCC spheroids (RCC7, 786-O and RCC10) in a hydrogel matrix that mimics the mechanical properties of tumor tissue and measured different invasive parameters such as maximum invasion length, invasion area, invasive cell counts, invasion speed and invasive trajectory. Over seven days, RCC7 spheroids exhibited the highest invasive behavior compared to 786-O and RCC10 spheroids. RCC10 spheroids remained compact, exhibiting minimal invasion and a directed, forward-biased migration pattern, possibly driven by weaker chemotactic cues or lower cellular plasticity ^47^. In contrast, RCC7 and 786-O cells exhibited erratic trajectories, which might reflect a more exploratory motility pattern. Their backward movements could imply cell-cell communication or “leader-follower” dynamics, where some cells may temporarily return to guide or recruit other cells ^48^. These dynamic movements align with their higher invasiveness, indicating an increased capacity to adapt to microenvironmental changes and exploit new invasion paths. Our findings suggest that trajectory complexity, rather than migration speed alone, may serve as a key indicator of tumor aggressiveness in our 3D invasion model.

Non-uniform distribution of genetic and phenotypic subpopulations within solid tumors causes many tumors of similar histological grade to have different metastatic potential, thus complicating existing prognostic assays ^49^. To further validate the relevance of our *in vitro* 3D model, we analyzed tumoroids derived from human ccRCC samples of different histopathological classifications. Consistent with their clinical grading, MC-156 tumoroids exhibited significantly higher invasion speed and exploratory motility than PD-157 tumoroids. While these results suggest a potential link between histopathological classification and invasive behavior, further studies with larger cohorts will be required to confirm this correlation robustly.

Our experiments highlight the limitations of 2D *in vitro* models in assessing RCC metastasis and the advantages of 3D spheroid models in capturing invasion dynamics and cell heterogeneity. Altogether, they suggest that testing cancer cell invasion in our 3D spheroid/tumoroid models may be predictive of *in vivo* tumor and metastatic progression, providing a promising platform for identifying high-risk patients and guiding treatment strategies. Future refinements, such as incorporating microfluidic chips to model the tumor microenvironment and metastatic cascade, could offer a powerful tool for studying tumor invasion and predicting therapeutic responses, paving the way for targeted therapies for metastatic RCC.

## Abbreviation

AVJ: arterio-venous junction
CAM: chorioallantoic membrane
ccRCC: clear cell Renal Cell Carcinoma
CTC: circulating tumor cells
CV: caudal vein
DA: dorsal aorta
EDD: embryonic development day
EMT: Epithelial-Mesenchymal Transition
hpf: hour post-fertilization
hpi: hour post-injection
MET: mesenchymal-to-epithelial
MMP2: matrix metalloproteinase 2
mRCC: metastatic Renal Cell Carcinoma
PD-1: receptor programmed death-1
PD-L1: Programmed death-ligand 1
PDX: Patient-Derived Xenografts
pEMT: partial Epithelial-Mesenchymal Transition
RCC: Renal Cell Carcinoma

## Author contributions

**Beatrice Cesana**: Conceptualization; formal analysis; investigation; visualization; writing – original draft. **Laurie Nemoz-Billet**: Investigation, writing – review and editing ; **Valentin Azemard** : Investigation, formal analysis, writing – review and editing ; **Catherine Pillet** : Investigation ; **Estelle Bigot**: Investigation ; **Nicolas Chaumontel** : Investigation ; **Jean-Luc Descotes** : Resources ; **Naël Osmani** : writing – review and editing ; **Jacky G. Goëtz** : writing – review and editing ; **Claude Cochet**: Conceptualization; supervision; investigation; writing – review and editing. **Odile Filhol**: Conceptualization; funding acquisition; supervision; project administration; writing – review and editing.

## Acknowledgment

We thank the animal unit staff (Bama S. Magallon C., Bailly S., Andrieux A. and Pointu H.) at Interdisciplinary Research Institute of Grenoble (IRIG) for animal husbandry and the PICSTRA imaging facility (CRBS). We sincerely thank Inovotion, Inc. (La Tronche, France) for their independent experiment of the tumor growth and metastasis analyses using the CAM assay, as well as their evaluation of chicken embryo toxicity. This work was supported by recurrent institutional funding from INSERM, CEA, and Ligue Contre le Cancer - Comité de l’Isère, Université Grenoble Alpes, Centre Hospitalier Universitaire de Grenoble-Alpes (CHUGA), Groupement des Entreprises Françaises dans la Lutte contre le Cancer (GEFLUC). VA is supported by PUI (Pôle Universitaire d’Innovation Grenoble). We thank GRAL LabEX, a program of the Chemistry Biology Health Graduate School of Université Grenoble Alpes (ANR-17-EURE-0003) (ANR-10-LABX-49-01).

Work and people supervised by JGG and NO are mostly supported by the INCa (Institut National Du Cancer, French National Cancer Institute), charities (La Ligue contre le Cancer, ARC (Association pour la Recherche contre le Cancer), FRM (Fondation pour la Recherche Médicale), Ruban Rose, Rohan Athlétisme Saverne and Trailers de la Rose), the National Plan Cancer initiative, the Region Grand Est, INSERM and the University of Strasbourg. This work has benefited from direct support by INCa (PLBIO23-255). LNB is supported by a post-doctoral fellowship from the La Fondation pour la Recherche Medicale.

## Conflict of interest statement

The authors declare no competing conflict of interests.

## Notes

### Competing Interest Statement

The authors have declared no competing interest.

