## supplemental figures and legends for "Using 3D Invasion properties of RCC Cell Lines *In Vitro* to predict their Metastatic Potential *In Vivo*"

**Supplementary Material and Methods**

### Preparation of cell extracts

RPTEC, ACHN, RCC10, 786-O and RCC7 cells were cultured to reach sub-confluency. Cells were washed with PBS and frozen at - 80°C. Cells were lysed on ice for 30 min in RIPA buffer (10 mM Tris-HCL, pH 7.4, 150 mM NaCl, 1% Triton X-100, 0.1% SDS, 0.5 % DOC, 1 mM EDTA) with protease and phosphatase inhibitor cocktail (Sigma Aldrich, P8340, P2850, P5726) at the recommended concentrations, centrifuged for 15 min at 4°C at 13.000 rpm and the supernatants collected. Proteins were quantified using the BCA protein assay kit (Pierce, ThermoFisher Scientific).

### Immunoblotting

Antibodies used for western blot analysis included GAPDH (#AM4300) from Invitrogen. HIF2$\alpha$ (#100-122) from Novus Biologicals. E-Cadherin (#610404), N-Cadherin (#610920), $\beta$1 Integrin (#610467 from BD Transduction Laboratories. Vimentin (#V5255), ZEB2 (#C83384) from Sigma Aldrich. PDL-1 (#13684), OCT4 (#2750), Snail1 (#3895), MMP2 (#4022), AKT (#9272), P-AKT S473 (#4060), P-Paxillin1 Y118 (#2541) from Cell Signaling Technology. Paxillin1 (#610052) from BD Biosciences. CXCR4 (#ab124824) from Abcam. After three washes, secondary antibodies (peroxidase-conjugated affinity pure anti-rabbit IgG (#111035OO3) or goat anti-mouse IgG (#115035003) from Jackson Immuno Research) were applied for 1 h followed by three more washes with TBST. Immobilon Forte Western HRP substrates (Millipore) was added and detection was achieved by using a Fusion FX acquisition system (Vilbert). Anti-GAPDH was used as a loading control and band intensities were quantified using ImageJ.

### 2D cell assays proliferation

786-O, RCC10, and RCC7 cells were seeded in 96-well plates (5,000/well). Cells were monitored for 5 days with CELLCYTE X™ ECHO (10X, every 2 h), cell proliferation was tracked in real time, and confluence over time (%/h) was analyzed via image software (Cellcyte Live Cell Analyzer software).

### 2D cell migration

786-O, RCC7, and RCC10 cells were seeded in 96-well plates (30,000 cells/well). After 24 h, mitomycin C was applied for 2 h to inhibit proliferation [16], and an 800 µm wound was created using WoundMaker (Essen Biosciences). Cells were monitored for 3 days with CELLCYTE X™ ECHO (10X, every 2 h), and confluence was analyzed over time (%/h).

### 2D cell invasion

786-O, RCC7 and RCC10 cells were seeded (200,000) cells in Matrigel-coated Boyden chambers (Corning® BioCoat Matrigel) with a 2 %-20 % FBS serum gradient. After 48 h at 37°C, cells were fixed in 4% PFA for 15 min and stained with Hoechst 33342 (1 µg/ml). Non-invasive cells were removed and membraned were mounted and observed (AxioObserver Z1, Zeiss, TILES mode, 10X). Invading cells were quantified using ImageJ.

#### Immunofluorescent labeling of tumoroids

Tumoroids were washed with PBS, fixed using 4% paraformaldehyde (PFA) (Sigma, 1 h, 4°C). All subsequent steps were carried out with gentle agitation (60 rpm). Following a 10 min PBS-Tween20 wash, samples were permeabilized and blocked for 1 h in Organoid Washing Buffer (OWB), containing 0.1% Triton-X100 and 0.2% BSA in PBS. Primary antibodies were incubated overnight (4°C), followed by 3 OWB washes (2 h) and secondary antibodies incubation overnight (4°C). On the third day, tumoroids underwent 3 additional OWB washes, with Hoechst (1 µg/ml) included in the second wash for nuclear staining. The primary antibodies used were CA9 (NovusBio, #NB100-417) and Phalloidin (Invitrogen, #A12379). The secondary antibody applied was Cy3 Goat anti-rabbit IgG (Jackson ImmunoResearch, #111-165-003). Imaging was performed using a Zeiss Apotome microscope.

#### Image analysis

To measure the average velocity of proliferation and wound closure over the measured time points (%/h), the confluence values were first normalized to account for variations in initial confluence across different cell types and replicates. The velocity was then calculated by performing linear regression analysis on the normalized confluence values.

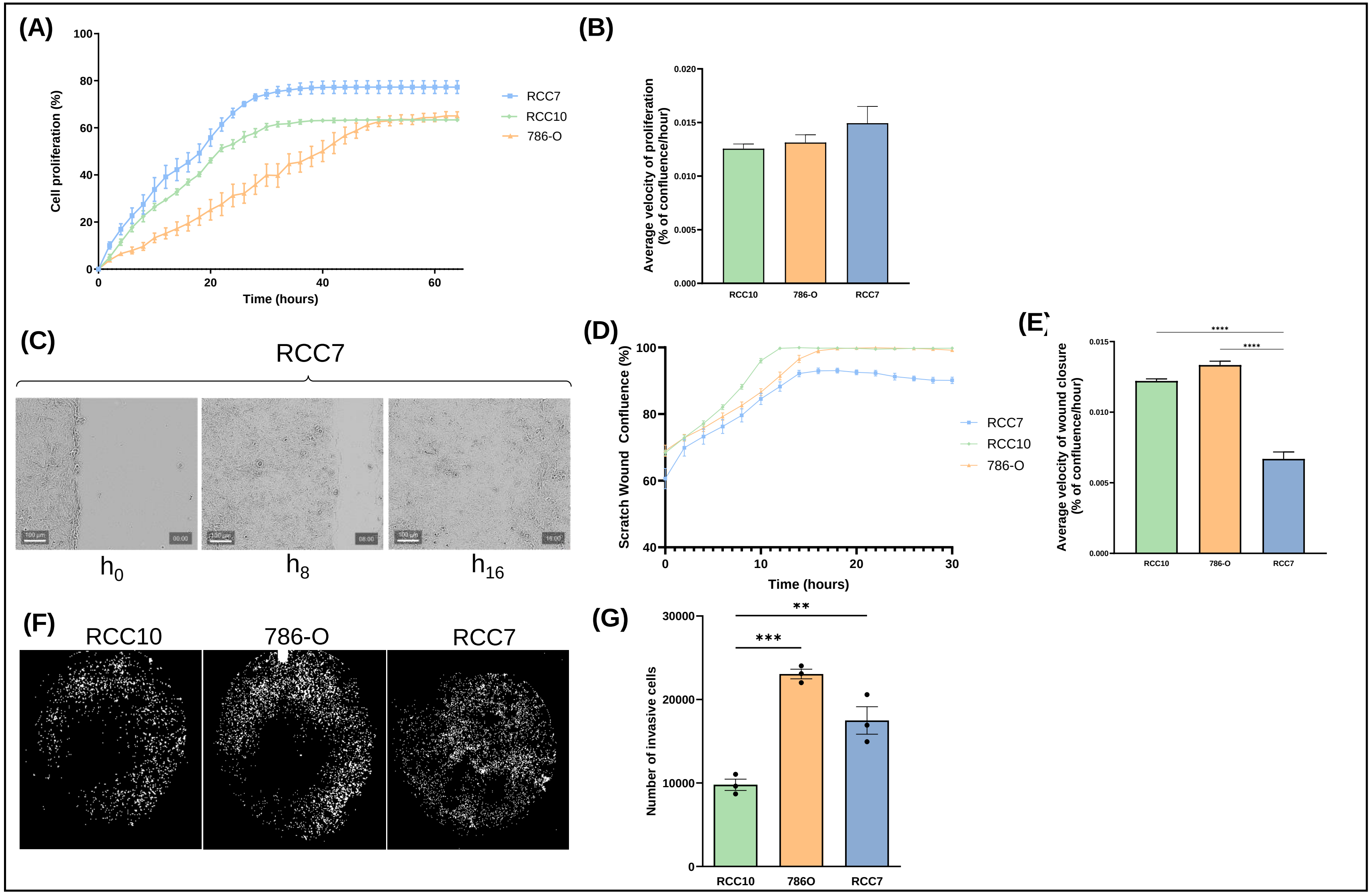


**Fig S1 : 2D Cell line-specific proliferation and migration (A)** Proliferation assay of RCC cell lines performed with CELLCYTE X™ Live Cell Imager And Analyzer **(B)** Average velocity of proliferation (% of confluence/hour) **(C)** Representation of the Scratch Wound Healing Assay for the RCC7 cell line **(D)** Scratch wound confluence data (%) for all RCC cells after 30 hours of culture, performed with CELLCYTE X™ Live Cell Imager. **(E)** Average velocity of wound closure (%of confluence/hour) **(F)** Image representation of the membranes from the Matrigel-coated Boyden invasion assays with indicated RCC cell lines. Cell nuclei are stained with Hoechst 33342. **(G)** Number of invasive cells counted on the bottom of each Matrigel-coated Boyden membrane. Data show mean ± SEM, with n = 6 for the proliferation assay, n = 12 for the Scratch Wound Healing Assay and n = 3 for the Matrigel-coated Boyden invasion assay. Significance was assessed using a one-way ANOVA with Tukey's multiple comparison test comparing each cell line to every other cell line. (***p*<0,01 ; ****p*<0,001 ; **** *p*<0,0001)

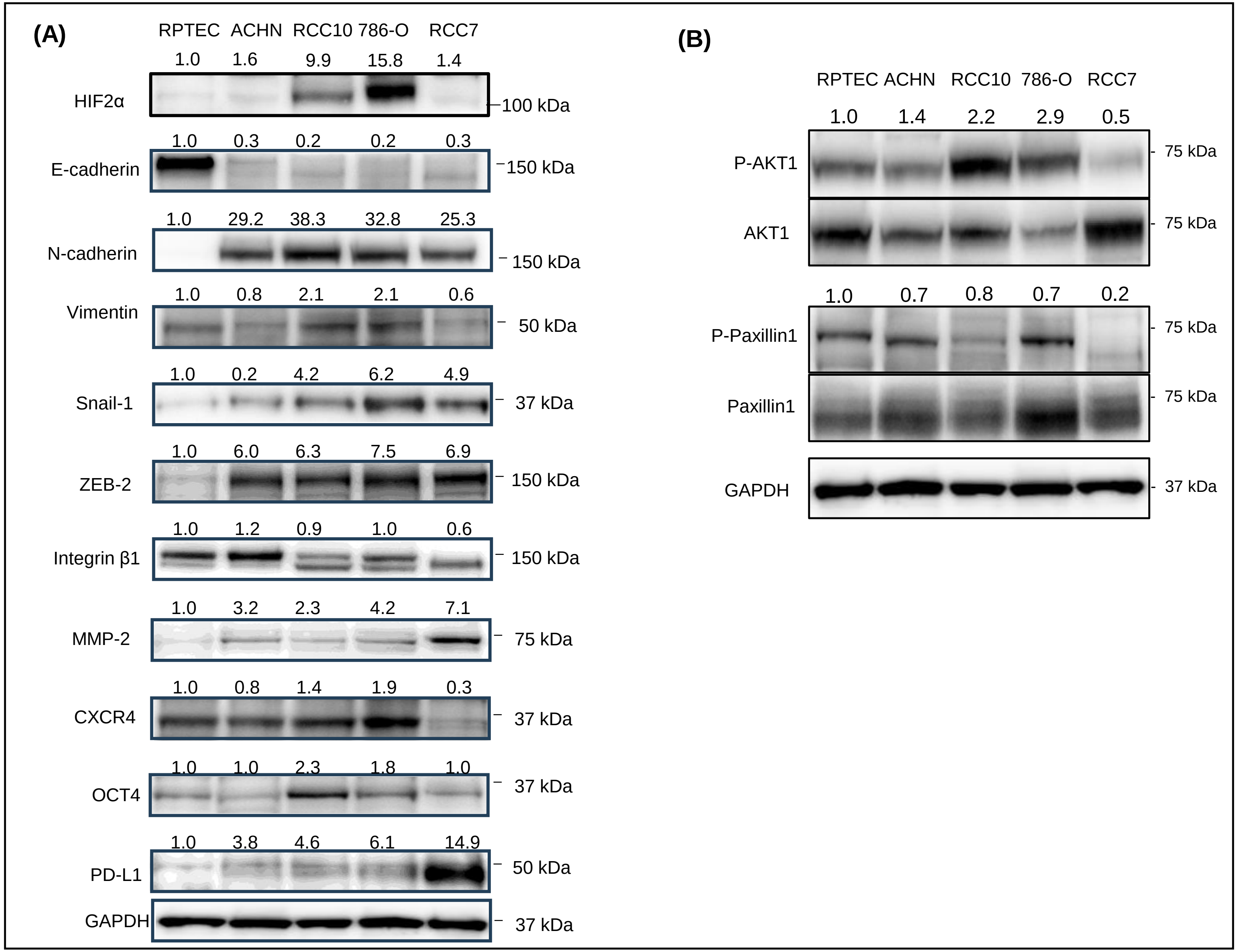


**Fig S2 : Western Blot analysis of key signaling pathways in RCC cell lines and in RPTEC, a healthy cell line.** Representative Western Blot analysis comparing proteins involved in the EMT **(A)** and phosphorylation pattern of AKT1 and Paxillin1 **(B).** GADPH was used for normalization of protein loading. Quantification of band intensities from the Western blot A and the ratio of the phosphorylated protein over the total protein B, are indicated above each lane.

(A)

| Mean Cq Alu Seq in lower CAM | | | |
|---|---|---|---|
| Samples | RCC10 | 786-O | RCC-7 |
| 1 | 25,73 | 25,095 | 24,625 |
| 2 | 23,535 | 20,905 | 23,985 |
| 3 | 24,19 | 23,08 | 23,805 |
| 4 | 24,23 | 24,11 | 24,005 |
| 5 | 24,41 | 21,67 | 23,415 |
| 6 | 24,285 | 24,14 | 23,72 |
| 7 | 24,315 | 23,995 | 23,81 |
| 8 | | | 23,425 |

| | |
|---|---|
| Mean Cq ALU seq CTRL | 24,405 |
| SD Alu seq CTRL | 0,2 |
| Background threshold | 24,205 |

(B)

| Mean Cq Alu Seq in lungs | | | |
|---|---|---|---|
| Samples | RCC10 | 786-O | RCC-7 |
| 1 | 28,14 | 22,06 | 27,14 |
| 2 | 23,93 | 26,96 | 25,89 |
| 3 | 27,01 | 24,00 | 25,95 |
| 4 | 26,71 | 27,55 | 26,50 |
| 5 | 27,35 | 26,47 | 26,67 |
| 6 | 27,48 | 26,99 | 24,89 |
| 7 | 24,90 | 27,04 | 26,61 |
| 8 | 27,34 | 27,16 | 27,54 |

| | |
|---|---|
| Mean Cq Alu seq CTRL | 26,71 |
| SD Cq Alu seq CTRL | 0,33 |
| Background threshold | 26,37 |

(C)

| Target Gene | Forward (5'-3') | Reverse (5'-3') | Probe (5'-3') |
|---|---|---|---|
| Alu sequences | GGTGAAACCCCGTCTCTACT | GGTTCAAGCGATTCTCCTGC | CGCCCGGCTAATTTTTGTAT |

(D)

| Gr. # | Group Description | Total | Alive | Dead | % Alive | % Dead |
|---|---|---|---|---|---|---|
| 1 | RCC10 | 17 | 16 | 1 | 94.12 | 5.88 |
| 2 | 786-O | 16 | 16 | 0 | 100 | 0 |
| 3 | RCC7 | 20 | 16 | 4 | 80 | 20 |

**Fig S3 : Quantitative Evaluation of Metastasis. (A)** Duplicates mean value of quantification cycles (Cq) measured by qPCR for human specific Alu sequences (Alu Seq) in the lower CAM. Green values show positive signal, determined by a mean Cq value higher than the background threshold of the qPCR corresponding with the mean value of Cq of control (CTRL) samples minus its standard deviation (SD). **(B)** Duplicates mean value of quantification cycles (Cq) measured by qPCR for human specific Alu sequences (Alu Seq) in the chicken lungs. Green values show positive signal. **(C)** Nucleotide sequences of forward, reverse and probe primers used for the detection of Alu sequences by qPCR based on the protocol described by Funakoshi & al [14]. **(D)** Number of dead and surviving embryos per group.
